# Germline-restricted chromosome of songbirds has different centromere compared to regular chromosomes

**DOI:** 10.1101/2025.01.23.634594

**Authors:** Jakub Rídl, Dmitrij Dedukh, Zuzana Halenková, Stephen A Schlebusch, Vladimír Beneš, Mireia Osuna Lopez, Tomasz S. Osiejuk, Francisco J Ruiz-Ruano, Alexander Suh, Tomáš Albrecht, Jiří Reif, Radka Reifová

## Abstract

Centromeres are an important part of chromosomes which direct chromosome segregation during cell division. Their modifications can therefore explain the unusual mitotic and meiotic behaviour of certain chromosomes, such as the germline-restricted chromosome (GRC) of songbirds. This chromosome is eliminated from somatic cells during early embryogenesis and later also from male germ cells during spermatogenesis. Although the mechanism of elimination is not yet known, it is possible that it involves a modification of the centromeric sequence on the GRC, resulting in problems with the attachment of this chromosome to the mitotic or meiotic spindle and its lagging during anaphase, which eventually leads to its elimination from the nucleus. However, the repetitive nature and rapid evolution of centromeres make their identification and comparative analysis across species and chromosomes challenging. Here, we used a combination of cytogenetic and genomic approaches to identify the centromeric sequence of two closely related songbird species, the common nightingale (*Luscinia megarhynchos*) and the thrush nightingale (*L. luscinia*). We found a 436-bp satellite repeat present in the centromeric regions of all regular chromosomes, making it a strong candidate for the centromeric repeat. This centromeric repeat was highly similar between the two nightingale species. Interestingly, hybridization of the probe to this satellite repeat on meiotic spreads suggested that this repeat is missing on the GRC. Our results indicate that the change of the centromeric sequence may underlie the unusual inheritance and programmed DNA elimination of the GRC in songbirds.

## Introduction

Centromeres are an essential component of chromosomes. By recruiting the kinetochore – a protein structure that mediates the attachment of the chromosomes to the microtubule spindle – they control chromosome segregation during cell division (McKinley and Cheeseman 2016). Defects in centromere function can lead to the loss or amplification of individual chromosomes in the dividing cells, with potentially deleterious consequences for the organisms (Holland and Cleveland 2009). However, alterations in the centromere have also been implicated in the unusual non-Mendelian inheritance (Clark and Akera 2021; Chmátal et al. 2017; Henikoff et al. 2001; Rosin and Mellone 2017) or programmed DNA elimination (Dedukh and Krasikova 2022) of certain chromosomes, including the parasitic B chromosomes (Johnson Pokorná and Reifová 2021).

Although the centromeric function is essential and well conserved, the centromeric DNA sequences, which are typically composed of satellite tandem repeats, represent one of the most rapidly evolving sequences in eukaryotic genomes (Henikoff et al. 2001; Poignet et al. 2021; Voleníková et al. 2023). This “centromeric paradox” can be explained by the frequent female meiotic drive occurring at centromeric loci, where centromeres with the ability to preferentially move the chromosome to oogonia rather than polar bodies can spread rapidly in the population and cause rapid turnover of centromeric sequences between species (Kursel and Malik 2018). In addition, centromeric sequences can evolve without much constraint, as epigenetic mechanisms leading to the loading of the centromere-specific histone H3 variant CENP-A, rather than the sequence itself, typically determine whether DNA acquires centromeric function (McKinley and Cheeseman 2016).

The rapid evolution of centromeric sequences, combined with their repetitive nature, makes them notoriously difficult to study. The position of the centromere within chromosomes can be inferred from cytogenetic data by observing primary constriction on metaphase chromosomes or by using various centromeric markers (Poignet et al. 2021; Talbert and Henikoff 2020). Furthermore, as genomic regions surrounding the centromere show reduced nucleotide diversity due to suppressed recombination, the centromeric position can be estimated from population genomic data (Begun and Aquadro 1992; Begun et al. 2007; Weissensteiner et al. 2017; Weighill et al. 2019). However, centromeric repeats themselves are often missing from genome assemblies and their identification requires a more detailed analysis of the repetitive genome content. As a result, centromeres have only been characterised in a limited number of species (e.g., Yamada et al. 2002; Yi et al. 2015; Khost et al. 2017; Weissensteiner et al. 2017; Uno et al. 2019; Blommaert et al. 2024; Chang et al. 2024). Only recently has the more detailed analysis of centromere repeat array structure been comprehensively possible with (nearly) gap-free genome assemblies for example in humans (Altemose et al. 2022; Gao et al. 2024; Logsdon et al. 2024) or chicken (Huang et al. 2023).

In songbirds, the most species-rich group of birds comprising nearly half of all bird species, the candidate centromeric repeats have only been described in few species, the chaffinch (*Fringilla coelebs*) (Saifitdinova et al. 2001), the hooded crow (*Corvus corone cornix*) (Weissensteiner et al. 2017), the zebra finch (*Taeniopygia guttata*) (Takki et al. 2022) and birds-of-paradise (Paradisaeidae species) (Peona et al. 2023). Songbirds have relatively conserved karyotypes consisting of a similar number of chromosomes across species (Kapusta and Suh, 2017). In addition to the standard set of chromosomes, songbirds contain the so-called germline-restricted chromosome (GRC) (Kinsella et al. 2019; Pigozzi and Solari 1998; Torgasheva et al. 2019). The GRC shows unusual mitotic as well as meiotic inheritance (Borodin et al. 2022). It is eliminated from somatic cells during early embryogenesis and from male germ cells during spermatogenesis (Pigozzi and Solari 2005, but see Pei et al. 2022). Although it is normally present as a single copy in males and two copies in females, polymorphism in copy number has been observed between individuals (Borodin et al. 2022; Malinovskaya et al. 2020; Torgasheva et al. 2021) and sometimes even between cells within an individual (Sotelo-Muñoz et al. 2022). Although the mechanisms behind this unusual inheritance are still unknown, it is possible that modification of the GRC centromeric sequence may be involved.

Here we used a multi-evidence approach to investigate the position and repetitive content of the centromeres in two closely related songbirds, the common nightingale (*Luscinia megarhynchos*) and the thrush nightingale (*L. luscinia*). The two species diverged approximately 1.8 Mya (Storchová et al. 2010) and still occasionally hybridise in the sympatric zone (Sottas et al. 2023). Recently a high-quality chromosome-level genome assembly has been published for both species (Schlebusch et al. 2023). First, we estimated the centromere position on individual chromosomes based on patterns of nucleotide diversity along the genome and previously published cytogenetic data (Poignet et al. 2021). Second, we searched for repetitive sequences corresponding in position to the estimated centromere positions from genomic and cytogenetic data. Finally, we generated probes for fluorescence in-situ hybridization (FISH) against the candidate repeats to cytogenetically verify that they are localised within the centromeric region.

We found a 436-bp satellite repeat present in the centromeric region of all regular chromosomes (i.e., autosomes and sex chromosomes) making it a good candidate for the centromeric sequence in nightingales. The cytogenetic data, however, suggest that the repeat is missing from the GRC. Our results indicate that the GRC may have a different centromere than the regular chromosomes, which could explain its unusual mitotic and meiotic behaviour.

## Materials and Methods

### Animals and sample collection

Four unrelated individuals (two females and two males) from each nightingale species were captured using mist nets and/or collapsible traps in allopatric populations to avoid sampling of interspecific hybrids. The common nightingales (*L. megarhynchos)* were captured in South-Western Poland by the Odra river, between the towns Wrocław (N 51.203°, E 16.965°) and Lubiąż (N 51.265°, E 16.470°). The thrush nightingales (*L. luscinia)* were captured in the vicinity of Lomża city (between N 52.970°, E 21.899°and N 53.134°, E 22.416°) in North-Eastern Poland. The birds were sampled in 2018 at the beginning of the breeding season between May 6 and May 15. Birds were euthanized by cervical dislocation. From female individuals heart samples were dissected, flash-frozen in liquid nitrogen and stored at -80°C for further DNA isolation and whole genome sequencing. From male individuals the testis and tibia were dissected for preparation of meiotic and mitotic spreads. The work was carried out in accordance with ethical animal research requirements of Poland according to Polish law (the Act On the Protection of Animals used for Scientific or Educational Purposes, 15.01.2015, item 266, implementing Directive 2010/63/EU of the European Parliament and of the European Council of 22.09.2010). Experiments on birds were approved by the General Directorate for Environmental Protection (permission no. DZP-WG.6401.03.123.2017.dl.3).

### Whole genome sequencing

DNA from heart tissue from each sampled female individual was isolated using MagAttract HMW DNA Kit (Qiagen) according to the manufacturer’s instructions. The DNA samples were sent to GeneCore (EMBL, Heidelberg) for standard Illumina paired-end sequencing using the HiSeq 2500.

In addition, we utilised previously published (Schlebusch et al. 2023) standard Illumina sequencing data from kidneys from two male individuals of each species (BioProject accession code PRJNA808609) and chromosome-level somatic genome assemblies together with respective Illumina data from one female individual of each species (BioProject accession codes PRJNA810511 and PRJNA810515).

### Estimates of nucleotide diversity along chromosomes

All the sequencing reads were quality trimmed using Trimmomatic v0.39 (Bolger et al. 2014). The trimmed Illumina reads from two female (this study) and two male (Schlebusch et al. 2023) individuals of each species were aligned to their respective reference genome (accessions GCA_034336665.1 and GCA_034336685.1) (Schlebusch et al. 2023) using bwa v0.7.17 (Li 2013). Samtools v1.14 (Danecek et al. 2021) was used to remove duplicate alignments with the fixmate and markdup functions. BCFtools’s mpileup and call commands (Danecek et al. 2021) were used for single nucleotide polymorphism (SNP) variant calling. SNPs with a quality score greater than 20 were selected using the varFilter command from VCFtools (Danecek et al. 2011). Nucleotide diversity (π) was calculated using VCFtools in 50 kb windows along individual chromosomes and visualised using the R (R Core Team 2023) package ggplot2 (Wickham 2016).

### Identification of candidate centromeric repeats

The Illumina reads from one female individual from each species (Schlebusch et al. 2023) were used for identification of repetitive elements using RepeatExplorer2 (Novák et al. 2010, 2013), including the TAREAN pipeline (Novák et al. 2017) in the Galaxy environment, both according to the author’s guidelines (Novák et al. 2020). From each species, 200 000 paired-end reads were randomly selected using seqtk v1.3-r106 (https://github.com/lh3/seqtk) and used separately for each species with RepeatExplorer2 with the cluster merging option. Consensus sequences of each resulting putative tandem repeat were duplicated to create dimers before being used with RepeatMasker v4.1.1 (Smit et al. 2015) to identify their location and abundance in each reference genome. The cumulative length of each satellite within 50 kb windows was calculated in R (R Core Team 2023) and plotted using ggplot2 (Wickham 2016).

BLASTN (Camacho et al. 2009) was used to calculate the sequence similarity between the consensus sequences of the candidate centromeric satellite found in both species (see Results). To identify possible homologues in other species, we used BLASTN search of the common nightingale candidate centromeric satellite against NCBInr database and also against previously identified repetitive elements in the collared flycatcher (*Ficedula albicollis*) (Suh et al. 2018).

The identified candidate centromeric satellites were inspected for the presence of the 17-bp CENP-B box motif by aligning (A/C)TTCGTTGGAAACGGGA, the proposed mammalian CENP-B box consensus sequence (Masumoto et al. 1989), using MAFFT v7.511 (Katoh et al. 2019).

### Preparation of mitotic and meiotic chromosome spreads

Mitotic metaphase chromosomes were prepared from bone marrow as has been described in Poignet et al. 2021. Briefly, after washing out bone marrow from the tibias of each bird, cells were cultivated in 5 ml of D-MEM medium (Sigma Aldrich), with the addition of 75 µl of colcemid (Roche) solution for 40 min at 37 °C. Afterward, D-MEM medium (Sigma Aldrich) was replaced by pre-warmed hypotonic 0.075 M KCl solution for 25 min at 37°C and then cells were fixed in 3:1 (methanol: acetic acid) fixative solution. After washing three times in a fixative solution (methanol:acetic acid, 3:1), tissues were stored at -20 °C until their use. To prepare chromosomal spreads, the cell suspension was dropped on a slide and the fixative solution was allowed to evaporate. The remaining chromosomal spreads were stained with Giemsa (5% Giemsa in 0.07 M phosphate buffer, pH 7.4).

Meiotic spreads of pachytene chromosomes were prepared from testes of reproductively active males following Peters et al. (1997). Testes were placed in hypotonic solution (30 mM Tris, 50 mM sucrose, 17 mM trisodium citrate dihydrate, and 5 mM EDTA; pH 8.2) and cut to release cells. After 50 min of incubation in hypotonic solution, testis tissue fragments were transferred to 200 µl of 100 mM sucrose and disaggregated. The resulting cell suspension (40 µl) was dropped onto a slide previously treated with 1% PFA and 0.15% of Triton X100 (Sigma Aldrich), and spread by slide inclination. Afterward, slides were placed in a humid chamber for 90 minutes, washed for 2 minutes in 1× PBS, and used for immunostaining. Pachytene chromosomes were immunostained with rabbit polyclonal anti-SYCP3 antibody (ab15093, Abcam), labelling the lateral elements of the synaptonemal complex (dilution 1:200) combined with human anticentromere serum (CREST, 15-234, Antibodies Incorporated) binding to centromere-specific proteins CENP-A, CENP-B, CENP-C (dilution 1:50) (Mchugh 2007) according to protocol described in Moens (2006). We used the following secondary antibodies: Alexa-594-conjugated goat anti-Rabbit IgG (H+L) (A32740, Invitrogen; dilution 1:200) and Alexa-488-conjugated goat anti-Human IgG (H+L) (A-11013, Invitrogen; dilution 1:200). Primary and secondary antibodies were diluted in PBT (3% BSA and 0.05% Tween 20 in 1× PBS) and incubated in a humid chamber for 90 min each. Slides were washed 3 times in 1× PBS and dehydrated through an ethanol row (50%, 70%, and 96%, 3 min each). Finally, all slides were dried and stained with DAPI in mounting medium Vectashield (Vector laboratories).

### Fluorescence *in situ* hybridization (FISH) of candidate centromeric repeats

We used FISH to cytogenetically verify that the candidate centromeric repeat (see Results) is located in the centromeric regions of chromosomes. We used two kinds of probes against its sequence. First, we used probes synthesised and labelled by PCR using the following primers (annealing temperature 61°C):

*L. luscinia:*

Forward 5’-GCTGTGCACACTTTCGCTTT-3’,

Reverse 5’-CCGTCTACCCCTCTCACAGA-3’;

*L. megarhynchos:*

Forward 5’-GAAGTGCCGTCTACCCCTCT-3’,

Reverse 5’-GTGAGATCTGTGGTGTGCTGT-3ߣ.

PCR probes were labelled with biotin-16-dUTP (Roche, Mannheim, Germany) or digoxigenin-11-dUTP (Roche, Mannheim, Germany). The primers were designed inside of the satellite monomer with the estimated PCR product length of 221 bp in *L. megarhynchos* and 199 bp in *L. luscinia*. However, since the satellite is organised in tandem repeats, the outcome of the PCR reaction is a mixture of the probes of different lengths (657 bp and 635 bp in *L. megarhynchos* and *L. luscinia*, respectively, if a satellite dimer is amplified, 1093 bp and 1071 bp if a trimer is amplified and so on). The obtained mixture of probes should thus cover the whole length of the satellite.

Second, we used the following commercially synthesised oligoprobes. The 20-bp probe for *L. luscinia* (5’-GCTGTGCACACTTTCGCTTT-3) was labelled with biotin-16-dUTP, and the 22-bp probe for *L. megarhynchos* (5’-GTTGTAGCAAAGCTGGTTCTGG-3’) was labelled with digoxigenin-11-dUTP (see Supplementary Figure 2 for the positions of PCR primers and oligoprobes in the satellite motif).

For FISH, chromosome slides were pretreated with 0.01% pepsin in 0.01 M HCl for 10 min, fixed with 2% paraformaldehyde for 10 min, washed in 1× PBS, dehydrated in an ethanol series (50%, 70% and 96%, 3 min each) and air-dried. In the case of pachytene chromosomal spreads, the pretreatment steps were omitted. The hybridization mixture for PCR probes contained 50% formamide, 10% dextran sulfate, 2× ЅЅС (saline-sodium citrate buffer), 200 ng (per slide) of either *L. luscinia* or *L. megarhynchos* probe, and 10-fold excess of salmon sperm DNA (Sigma-Aldrich) to the amount of the probe. For two-colored FISH, we added 100 ng of *L. luscinia* probe labelled with biotin and 100 ng of *L. megarhynchos* probe labelled with digoxigenin to the hybridization mixture which was otherwise the same as previously described. In the case of oligoprobes, the hybridization mixture included 40% formamide, 10% dextran sulfate, 2× ЅЅС, 200 ng of oligoprobe, and a 10-fold excess of salmon sperm DNA (Sigma-Aldrich) to the amount of the probe. Both PCR and oligoprobes were denatured at 86°C for 6 minutes and placed on ice for 10 minutes. In the case of two-coloured FISH, the denatured probes were also incubated at 37 °C for 30 minutes. Simultaneously, chromosomal DNA on slides with mitotic chromosomes and pachytene chromosomes was denatured at 76 °C for 3 min, dehydrated in a series of ice-cold ethanol washes (50%, 70%, and 96%, 3 min each), and air dried. Afterward, a hybridization mixture with denatured probe was dropped on slides, covered by cover slides, mounted with rubber cement, and incubated overnight at either 37 °C in the case of PCR probes or at room temperature in the case of oligoprobes. After hybridization overnight, slides with PCR probes were washed thrice in 0.2× SSC for 5 min at 50 °C; slides with oligoprobes were washed in 2× SSC for 5 min at 46 °C. Detection of probes was performed with streptavidin conjugated with Alexa 488 (S11223, Invitrogen, San Diego, CA, United States); digoxigenin was detected by anti-digoxigenin antibodies conjugated with rhodamine (11207750910, Merck, Darmstadt, Germany). After incubation for 3 hours, slides were washed in 4× SSC for 5 min at 45°C, dehydrated in an ethanol series (50%, 70%, and 96%, 3 min each), and mounted with Vectashield/DAPI (1.5 mg/mL) antifade medium (Vector Laboratories, Burlingame, CA, USA).

Since we did not observe any difference between the mapping of PCR probes and oligoprobes, only results from PCR probes are shown in the results section.

### Image Processing

Chromosomal slides were examined by an Olympus Provis AX 70 epifluorescence microscope and ZEISS Axio Imager.Z2 epifluorescence microscope. Images of metaphase chromosomes were captured with an Olympus DP30BW CCD camera and a CoolCube 1 camera (MetaSystems, Altlussheim, Germany). The IKAROS and ISIS imaging software (Metasystems, Altlussheim, Germany) were used to analyse images adjusted in Adobe Photoshop software, version CS5.

## Results

### Estimation of centromere position within individual chromosomes

To estimate the position of the centromere on individual chromosomes, we first utilised whole genome sequence data from four individuals per species and calculated nucleotide diversity (π) along individual chromosomes in 50 kb windows. Figure 1 shows the distribution of π along the first four largest chromosomes (data for other chromosomes are shown in Supplementary Figure 1). In most chromosomes we found a single prominent region with reduced π values compared to the rest of the chromosome, representing a candidate centromeric region (Figure 1, Supplementary Figure 1).

**Figure 1.**
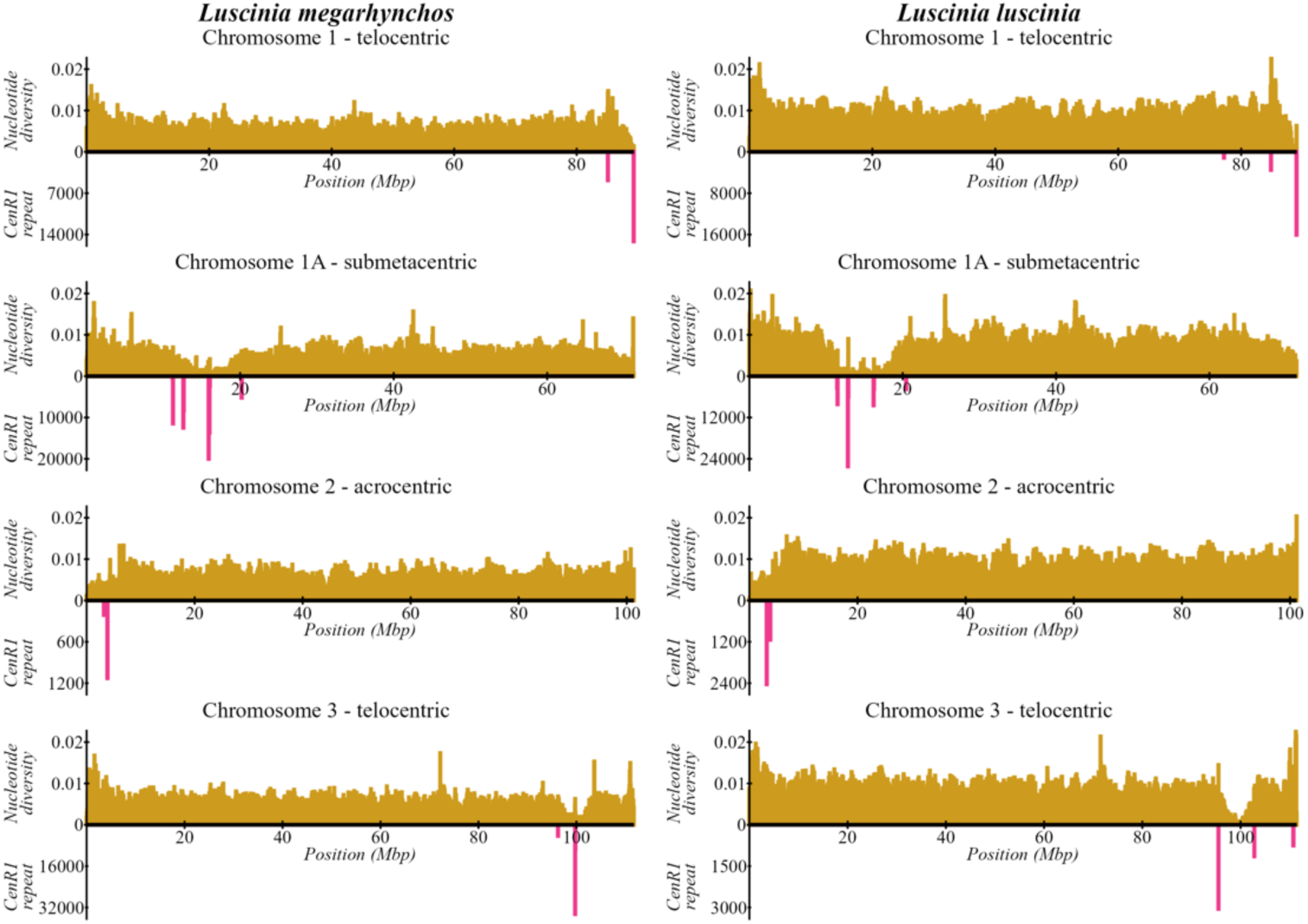
Nucleotide diversity (yellow) and the cumulative length of centromeric satellites *LmegCenR1* and *LlusCenR1* (magenta) in 50kb sliding windows along four largest nightingale chromosomes. Chromosome morphology classifications adapted from Poignet et al. (2021).

We then used previously published cytogenetic data (Poignet et al. 2021), where centromeres on pachytene chromosomes were stained with the CREST antibody. The centromere position on each chromosome was estimated by comparing the relative length of the short and long chromosome arms and all chromosomes were classified as telocentric, acrocentric, submetacentric or metacentric based on the estimated centromere position. The classification of the chromosomes based on the cytogenetic data corresponded with the position of the predicted centromeric regions based on the genomic regions of reduced genetic diversity for all chromosomes, where we were able to homologize the assembled chromosomes with cytogenetic data based on the chromosome lengths (Figure 1, Supplementary Figure 1).

The final estimated position of the centromeres on individual chromosomes was the same for both nightingale species, suggesting that no structural changes affecting the centromere position occurred after the species’ divergence.

### Identification of candidate centromeric repeat sequences

RepeatExplorer2 was used to identify repetitive sequences in both nightingale genomes, while RepeatMasker was used to identify the locations and abundance of these repeat sequences in the genomes. We found 5 tandem repeats in the common nightingale and 8 tandem repeats in the thrush nightingale (Supplementary Table 1 and 2). The most frequent repeat in both species was a 436-bp satellite, which was present in 6.4% of the analysed reads in *L. megarhynchos* and in 4.0% of the analysed reads in *L. luscinia*. The genome localization of the identified repeats was further compared to the estimated position of the centromere based on nucleotide diversity and cytogenetic data. The most abundant tandem repeat was the only repeat whose position corresponded with the predicted centromeric regions (Figure 1, Supplementary Figure 1). It was found almost exclusively in centromeric regions and was missing in other parts of the genome. None of the other putative repeats showed such a clear co-localization across chromosomes. This makes the most common tandem repeat the best candidate for the first identified centromeric repeat in nightingales, hence we refer to it as CenR1 in both species and name it as *LmegCenR1* in the common and *LlusCenR1* in the thrush nightingale.

The overall sequence identity between the CenR1 consensus sequences of the two nightingale species was 98.4% (Supplementary Figure 2), showing the high similarity of this candidate centromeric repeat in the two nightingale species. The *LmegCenR1* sequence of *L. megarhynchos* also showed 81% identity to the previously identified 436-bp satellite repeat (*fAlbSat4*) in the genome of collared flycatcher (*Ficedula albicollis*) (Suh et al. 2018). In addition, nucleotide blast search against NCBInr database showed hits to chromosomes of European robin (*Erithacus rubecula*, GenBank assembly accession GCA_903797595.2) with average identity 78%.

In humans, a 17-bp long motif called CENP-B box was identified in centromeric satellites as a binding site for centromere protein B which plays a significant role in centromere assembly (Masumoto et al. 1989). Putative CENP-B box with sequence similarity to human consensus sequence was later identified in other species (Gamba and Fachinetti 2020) including several bird taxa, e.g. pigeon, chaffinch and chicken (Solovei et al. 1996; Krasikova et al. 2012). We identified a putative CENP-B box motif in the CenR1 satellite in both nightingale species sharing the same sequence with overall 65% similarity to the mammalian CENP-B box consensus (Supplementary Figure 3).

### Candidate centromeric repeat co-localizes with centromeric markers

To further verify that the CenR1 represents the centromeric repeat, we prepared probes against these satellites in both nightingale species and hybridised them on mitotic and meiotic chromosome spreads of the corresponding species. In mitotic spreads, the signals were observed in centromeric regions, visible as chromosomal primary constrictions, in all macrochromosomes as well as microchromosomes. The intensity of the signals, nevertheless, varied in size among the chromosomes in both species (Supplementary Figure 4), suggesting that individual chromosomes may differ in the copy number of satellite repeats. A similar pattern was observed on pachytene spreads, where synaptonemal complexes were labelled with antibodies against their lateral components (SYCP3) and the centromeres were stained using the CREST antibody. We observed a complete overlap of signals between the CREST antibodies and CenR1 repeat in all regular chromosomes of both species (Figure 2), strongly suggesting that this satellite represents the centromeric repeat on all regular chromosomes.

**Figure 2.**
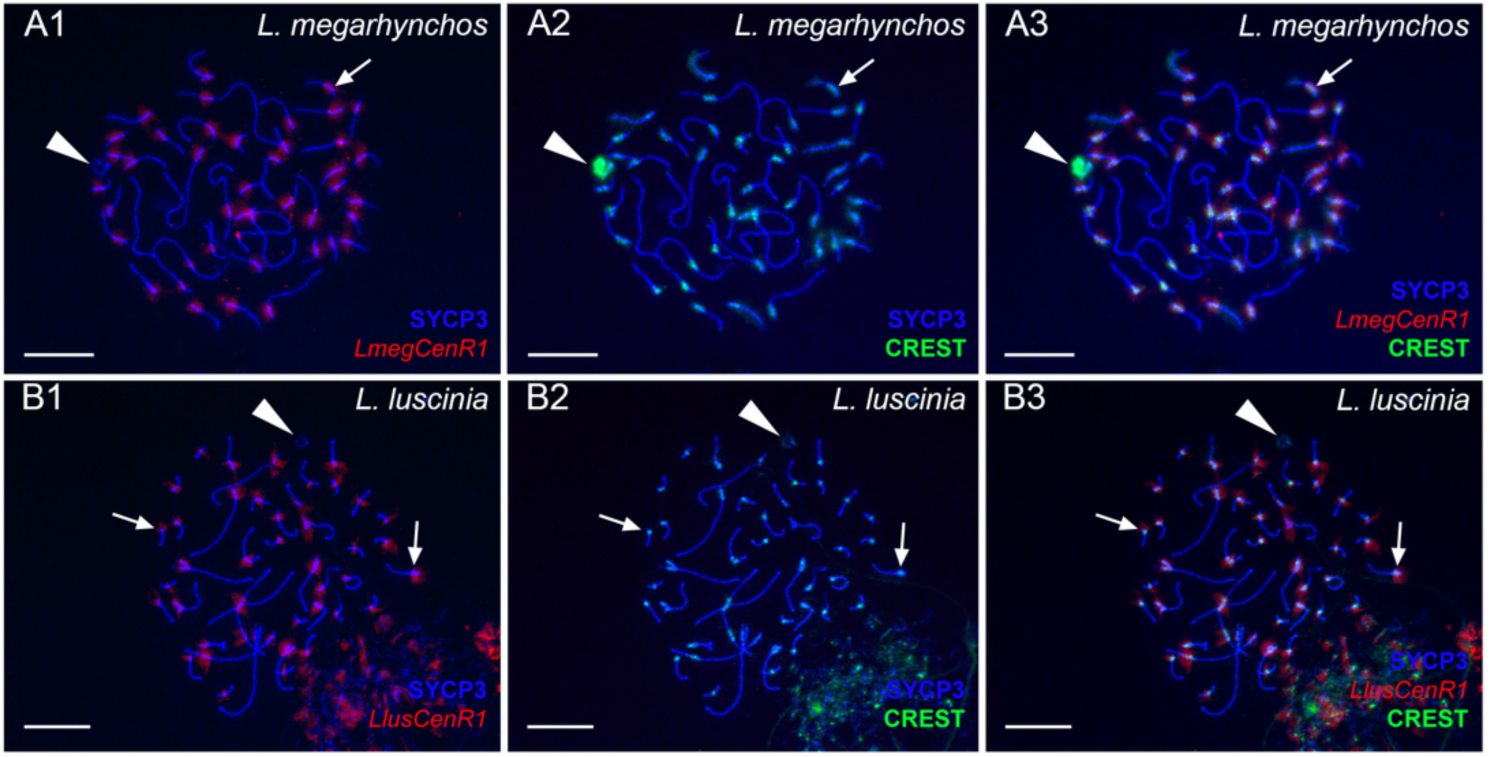
Simultaneous detection of centromeric regions by anti-centromere antibodies and *LmegCenR1* repeat on *L. megarhynchos* SC spreads (A1-A3) and *LlusCenR1* repeat on *L. luscinia* SC spreads (B1-B3). Lateral components of synaptonemal complexes were stained by anti-SYCP3 antibodies (blue in all panels). FISH-mapping of pericentromeric repeats (red in A1, B1, and A3, B3) as well as antibodies-based detection by CREST antibodies (green in A2, B2, and A3, B3) showed their colocalization (A3, B3) in all chromosomes (indicated by arrows) except GRCs (indicated by arrowheads). GRCs, represented as univalent in both species (indicated by arrowheads), exhibit CREST staining in the pericentromeric region and diffuse staining of its chromatin but do not show the signal from *LmegCenR1* and *LlusCenR1* repeats. Scale bar = 10 µm.

We further performed a cross-species experiment and hybridised the *L. megarhynchos* CenR1 PCR probe to *L. luscinia* chromosomes and vice versa. Hybridization was effective in both directions (Supplementary Figure 5A, B), which is consistent with the finding that the CenR1 satellite shows high similarity between the nightingale species. The two-coloured FISH experiment also showed perfect colocalization of the PCR probes of the two nightingale species in the centromeric regions on spreads of both species (Supplementary Figure 5C, D).

### GRCs have different centromeric sequences than other chromosomes

We identified the GRC in pachytene spreads from testis in both nightingale species. The GRC exists as a small univalent and unlike other chromosomes is labelled diffusely along its entire length by the CREST antibody (Poignet et al. 2021; Sotelo-Muñoz et al. 2022; Torgasheva et al. 2019). Interestingly, in contrast to other chromosomes, the CenR1 probe showed no signal on the GRC in either of the two nightingale species (Figure 2A1-B3), suggesting that the CenR1 repeat is missing, has very low abundance or is modified on this chromosome.

## Discussion

Centromeres play a crucial role in chromosome segregation during cell division through their ability to attach chromosomes to the mitotic or meiotic spindle. However, although high-quality reference genome assemblies now exist for many species, centromeric sequences are often missing or unidentified in genome assemblies due to their repetitive content. Here we have identified a candidate centromeric sequence in two closely related nightingale species. It consists of a 436-bp satellite repeat and is present in centromeric regions of all chromosomes except the GRC. Below we discuss our findings in the context of centromere evolution in birds and consider the possible implications of the different centromeric sequence for the inheritance of the GRC.

In birds, centromere composition has so far only been well described in the chicken. In this species, most acrocentric chromosomes have centromeres that are composed of a tandem array of the same 41-bp repeat called CNM (Huang et al. 2023). Although the CNM repeat appears to be an important part of the centromeres, the kinetochore itself does not usually bind to its sequence, but to its boundaries or to a smaller variable repeat motif embedded within the CNM (Huang et al. 2023). On the other hand, centromeres of metacentric or submetacentric chromosomes consist of chromosome-specific tandem repeats (Piégu et al. 2018).

Among songbirds, centromeric repeats have been identified in the zebra finch (Takki et al. 2022), chaffinch (Saifitdinova et al. 2001), and Corvoidea (crows, birds-of-paradise and relatives; Weissensteiner et al. 2017; Peona et al. 2023). In the zebra finch, analysis of the repetitive content of the genome, followed by FISH with the candidate repeats on the metaphase chromosomes, revealed that the centromeres of all chromosomes are composed of two distinct tandem repeats, *Tgut191A* and *Tgut716A*, with monomer lengths of 191 bp and 716 bp, respectively (Takki et al. 2022). In the chaffinch, genome restriction digestion followed by cloning of the repeat sequence and verification of the centromeric location by FISH led to the identification of a 505-506-bp tandem repeat present in the centromeric regions of all chromosomes (Saifitdinova et al. 2001). In Corvoidea, the combination of tandem repeat identification across species with population genomic data of hooded crows identified candidate centromeric repeats, pericentromeric repeats, and centromere-containing assembly gaps across autosomes (Weissensteiner et al. 2017; Peona et al. 2023). Here we identified the centromeric tandem repeat, called CenR1, with a monomer length of 436 bp in two other songbird species, the common nightingale and the thrush nightingale. While these two closely related species have the same tandem repeat in their centromeres, it shows no similarity to the tandem repeats identified in zebra finch and chaffinch. However, we have identified the homologous tandem repeat to CenR1 in the genomes of the collared flycatcher, previously described as *fAlbSat4* (Suh et al. 2018), and the European robin. These two species, together with nightingales, belong to the family Muscicapidae (Zhao et al. 2023). Therefore, it is possible that this centromeric sequence represents an ancestral centromeric sequence in this taxon, although this needs to be verified.

The centromeric function of the CenR1 satellite repeat in nightingales is supported not only by its presence in the centromeric regions of all regular chromosomes, as has been verified by the FISH analysis, but also by the presence of the CENP-B box in its sequence. The CENP-B box is a 17-bp sequence motif that is recognized by the kinetochore-associated protein CENP-B and is common in centromeric satellite sequences of many species (Gamba and Fachinetti 2020; Saffery et al. 1999). Moreover, a probe against the CenR1 tandem repeat co-localizes with the CREST anticentromere serum, which is known to bind the kinetochore proteins CENP-A, CENP-B and CENP-C (Mchugh 2007). However, to prove that the kinetochore binds directly to the CenR1 satellite repeat and not to a small embedded or nearby sequence, an antibody to the CENP-A kinetochore protein would need to be developed and used to immunoprecipitate the DNA directly bound to it, as has been done in chicken (Shang et al. 2010, 2013). Even if the CenR1 satellite does not form a core of the centromere and does not bind directly to the kinetochore, it can form a pericentric heterochromatin, which is important for sister chromatin cohesion and accurate chromosome segregation.

Interestingly, the GRC was the only chromosome that showed no signal in the FISH experiment with the CenR1 probe, suggesting that this repeat is either missing on the GRC, or has a lower number of copies than can be detected by FISH, or has diverged sufficiently so that probe does not bind to it. Unfortunately, the currently available GRC assembly is highly fragmented and still incomplete (Schlebusch et al. 2023), preventing us from analysing the repetitive content of the GRC and thus identifying the centromeric repeat on this chromosome. Notably, the GRC also shows different staining with the CREST serum compared to the regular chromosomes. While CREST only stains the centromeric regions on regular chromosomes, it weakly labels the entire length of the GRC in male pachytene cells. This may indicate that the kinetochore binds to the GRC in a different way compared to regular chromosomes, which might be related to its unusual behaviour in male meiosis.

In several systems, it has been shown that the inability of chromosomes to attach to the mitotic/meiotic spindle or the failure of chromatid separation and their subsequent lagging in the anaphase can lead to their elimination from the nucleus (Dedukh and Krasikova 2022; Ishii et al. 2016). We therefore propose that a modification of the centromeric sequence of the GRC could be the underlying mechanism for the programmed elimination of this chromosome from somatic cells and male germ cells. The problems with chromatid separation and chromosome lagging in anaphase can also lead to the duplication of the chromosome in one daughter cell while it is eliminated from the other cell if cell division is asymmetric (Banaei-Moghaddam et al. 2012; Johnson Pokorná and Reifová 2021; Ruban et al. 2020; Wu et al. 2019). Modification of the GRC centromere could thus also explain how the chromosome is duplicated in the female germline at some point during germline development and how the GRC copy number variation occasionally arises. Further cytogenetic analyses of early embryos and male and female gonads are required to test this hypothesis.

## Acknowledgments

This research was funded by the Czech Science Foundation (grant 23-07287S to R.R., T.A. and D.D.), the Grant Agency of Charles University (grant 314222 to Z.H.) and the Institutional Research Support grant of the Charles University (SVV 260684/2024).

## Author contributions

The project was conceptualised by R.R., and J.Rí.; Samples were collected by J.Re., T.A. and T.S.O.; Cytogenetic analysis was performed by D.D.; DNA extraction was done by J.Rí. and Illumina sequencing was done by M.O.L. and V.B.; Bioinformatic analyses were done by J.Rí., Z.H., and S.A.S. with consultation from A.S. and F.J.R-R; Manuscript was written by J.Rí., R.R., D.D., Z.H. and S.A.S and all authors contributed to text editing.

## Data archiving

The Illumina whole genome sequencing data generated in this study have been deposited in the NCBI’s SRA database under the BioProject accession code (PRJNA1178309). The consensus sequences of *LmegCenR1* and *LlusCenR1* have been deposited in the NCBI’s GenBank database (accession numbers PQ530385 and PQ530386, respectively).

## Research Ethics Statement

The work was carried out in accordance with ethical animal research requirements of Poland according to Polish law (the Act On the Protection of Animals used for Scientific or Educational Purposes, 15.01.2015, item 266, implementing Directive 2010/63/EU of the European Parliament and of the European Council of 22.09.2010). Experiments on birds were approved by the General Directorate for Environmental Protection (permission no. DZP-WG.6401.03.123.2017.dl.3).

