## Supplementary figures for "Germline-restricted chromosome of songbirds has different centromere compared to regular chromosomes"

**The document includes:**

Supplementary Figure 1
Supplementary Figure 2
Supplementary Figure 3
Supplementary Figure 4
Supplementary Figure 5
References

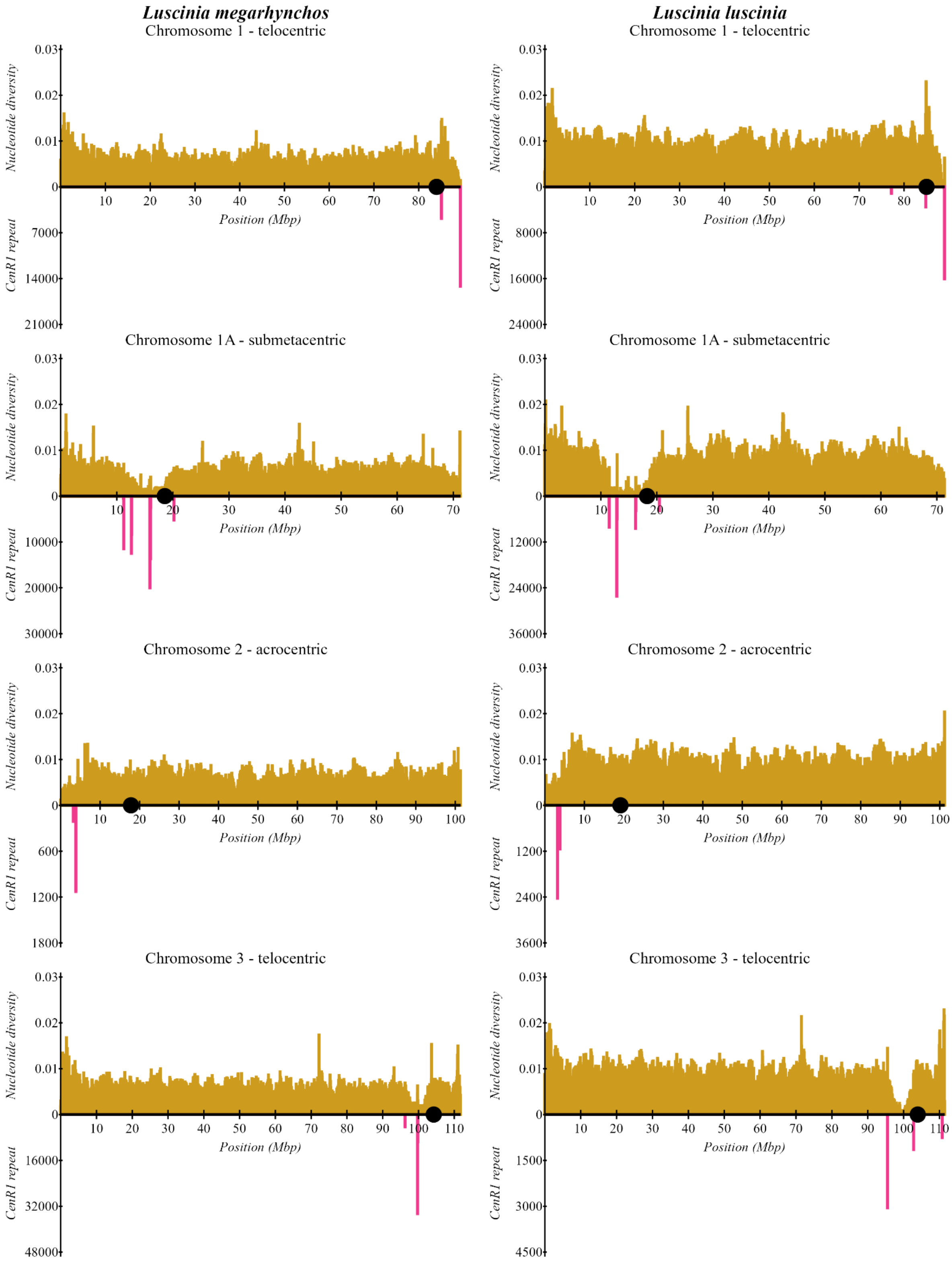

Luscinia megarhynchos
Luscinia luscinia
Chromosome 1 - telocentric
Chromosome 1A - submetacentric
Chromosome 2 - acrocentric
Chromosome 3 - telocentric
Nucleotide diversity
CenR1 repeat
Position (Mbp)
0.03
0.02
0.01
0
7000
14000
21000
8000
16000
24000
10000
20000
30000
12000
24000
36000
600
1200
1800
2400
3600
16000
32000
48000
1500
3000
4500

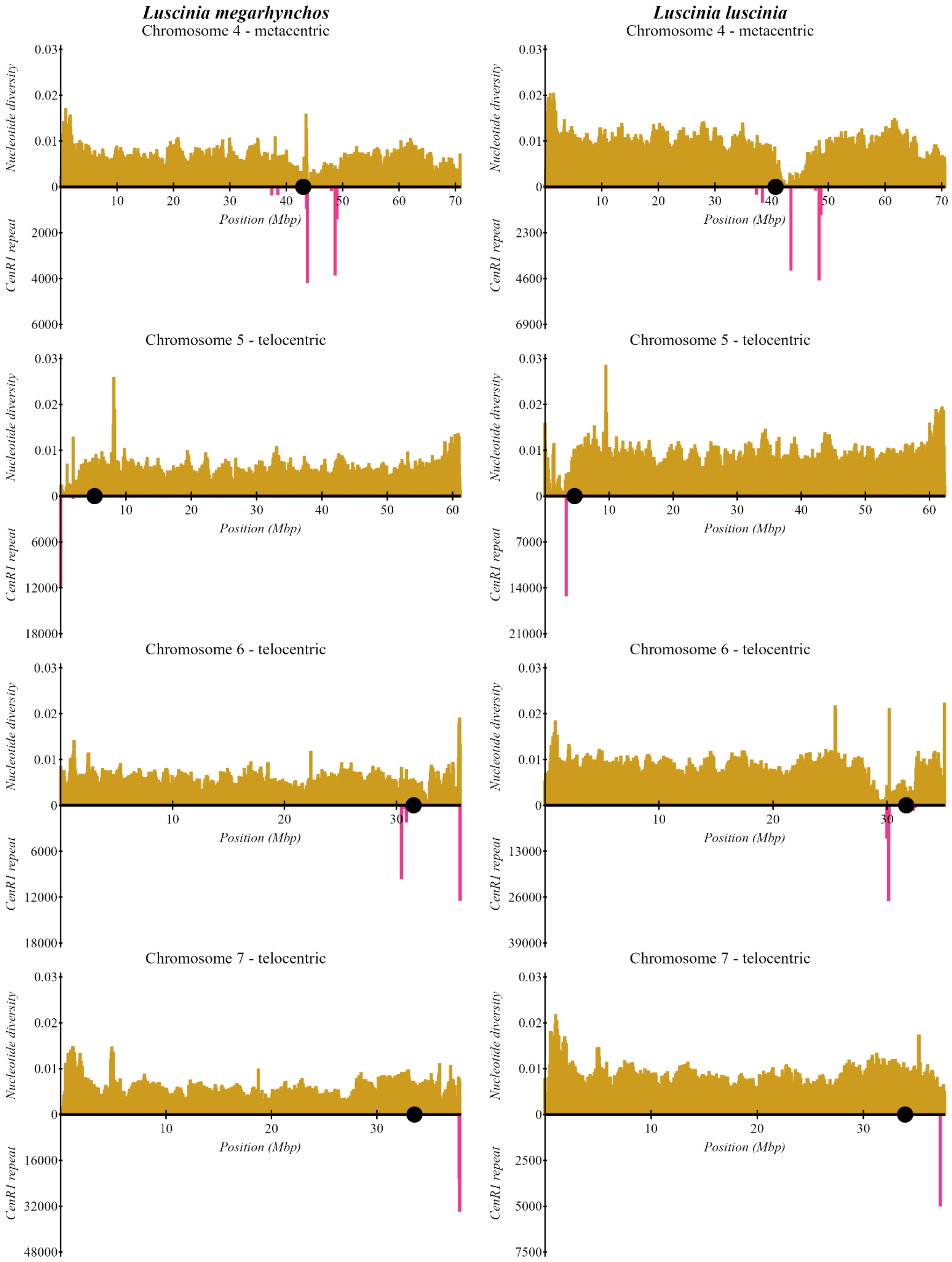

Luscinia megarhynchos
Chromosome 4 - metacentric
Luscinia luscinia
Chromosome 4 - metacentric
Chromosome 5 - telocentric
Chromosome 6 - telocentric
Chromosome 7 - telocentric
Nucleotide diversity
CenR1 repeat
Position (Mbp)
0.03
0.02
0.01
0
10
20
30
40
50
60
70
2000
4000
6000
2300
4600
6900
12000
18000
7000
14000
21000
13000
26000
39000
16000
32000
48000
2500
5000
7500

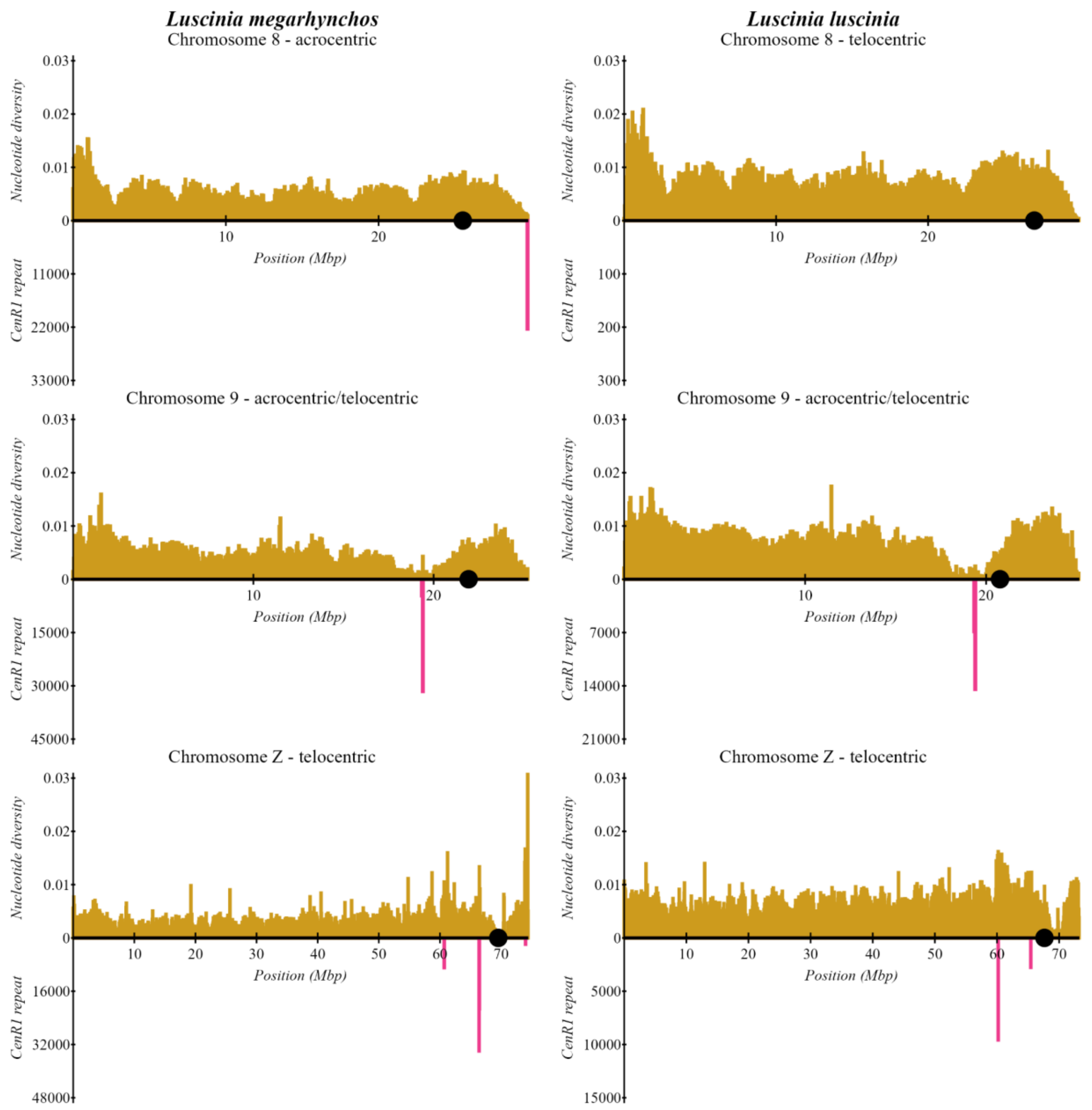


**Supplementary Figure 1.** Nucleotide diversity (π) (yellow) and the cumulative length of the centromeric satellites *LmegCenR1* and *LlusCenR1* (magenta) in 50kb sliding windows along the nightingale chromosomes. The ten largest chromosomes and the Z chromosome, which we were able to homologize with the karyotype data (Poignet et al. 2021) based on the chromosome lengths are shown. Chromosome morphology classification (telocentric, acrocentric, submetacentric and metacentric) is based on the estimated position of centromere from cytogenetic data. This position shows the median arm ratio (Poignet et al. 2021) and is marked by black dots on individual chromosomes. The estimated positions of centromeres from cytogenetic data roughly colocalize with the genomic regions of decreased nucleotide diversity and with the regions containing CenR1 repeats, although some discrepancies are present, likely caused by missing parts of our assemblies. The absence of CenR1 repeats in the *Luscinia luscinia* assembled chromosome 8 is likely a consequence of an unassembled end of this chromosome. This is supported by the location of the CenR1 peak in *Luscinia megarhynchos* at the very right end of the chromosome.

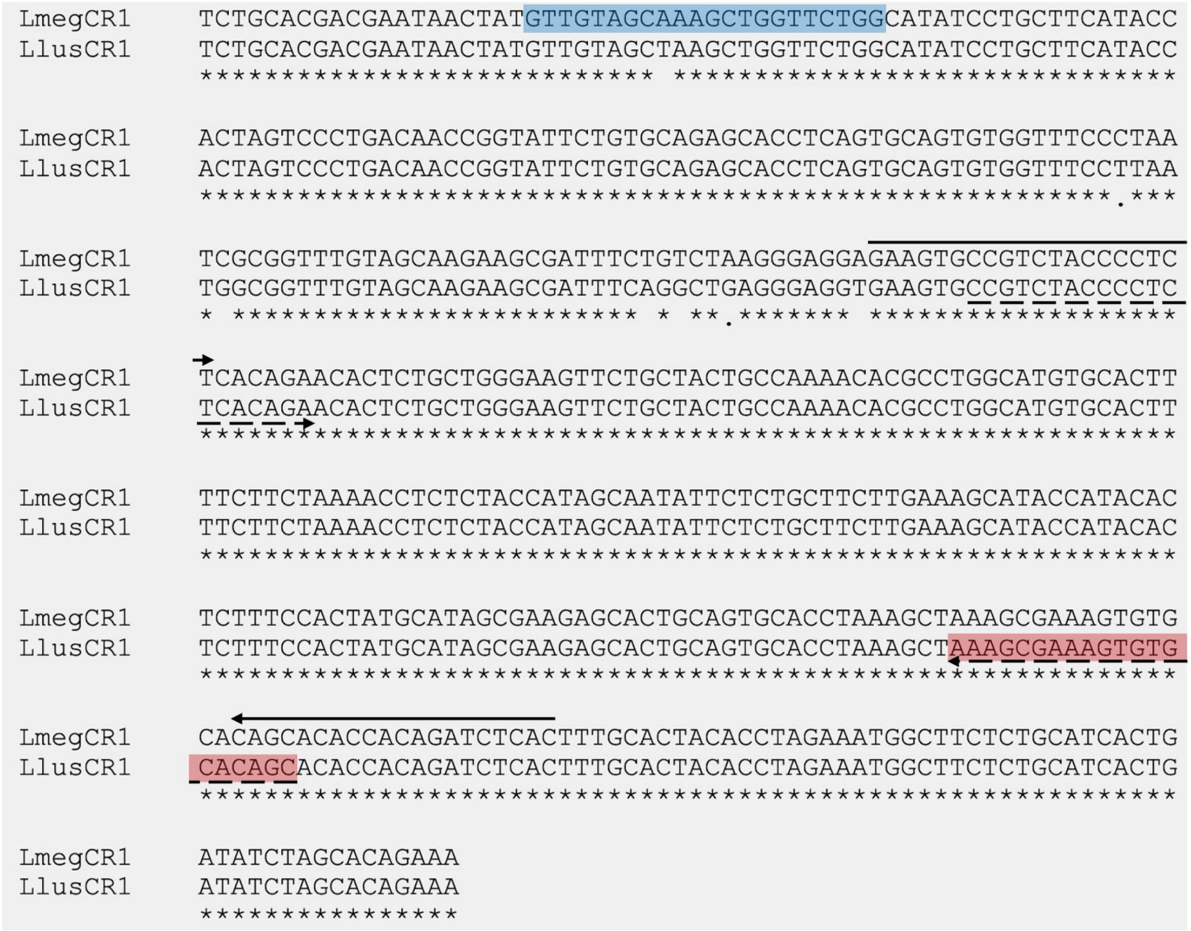


**Supplementary Figure 2.** Visualization of the *LmegCenR1* and *LlusCenR1* consensus sequence alignment using MAFFT v7.511 (Katoh et al. 2019). The position of the PCR primers used to generate the PCR probes are marked above the sequences by black arrows for *L. megarhynchos* and below the sequences by dashed arrows for *L. luscinia*. Sites targeted by oligoprobes are indicated in blue for *L. megarhynchos* and in red for *L. luscinia*.

```
                          ****    *  ****         identity
Consensus            (A/C)TTCGTTGGAAACGGGA
LmegCenR1rc              AGTGGTATGAAGCAGGA              65%
LlusCenR1rc              AGTGGTATGAAGCAGGA              65%
```

**Supplementary Figure 3.** Identification of a putative 17bp CENP-B box sequence in the *LmegCenR1* and *LlusCenR1* centromeric satellites with similarity to the mammalian consensus sequence. Nucleotides necessary for CENP-B binding in humans (Ohzeki et al. 2002) are marked by asterisks, nucleotides identical to the consensus are indicated in red, rc indicates reverse complementary sequence.

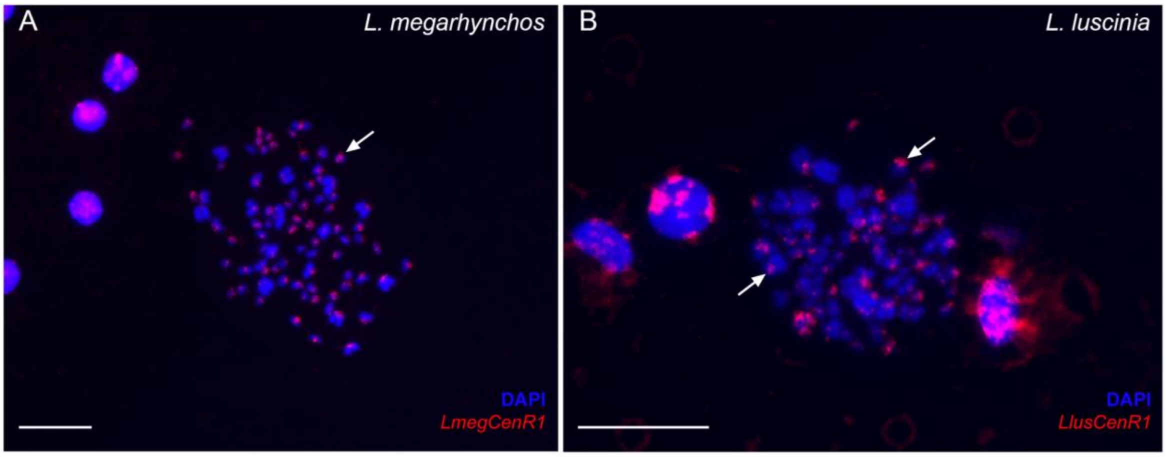


**Supplementary Figure 4.** FISH-based mapping of *LmegCenR1* repeat on *L. megarhynchos* mitotic chromosomes (A) and *LlusCenR1* repeat on *L. luscinia* mitotic chromosomes (B). Signals localise in centromeric regions of all chromosomes (indicated by arrows). Chromatin stained with DAPI. Scale bar = 10 µm.

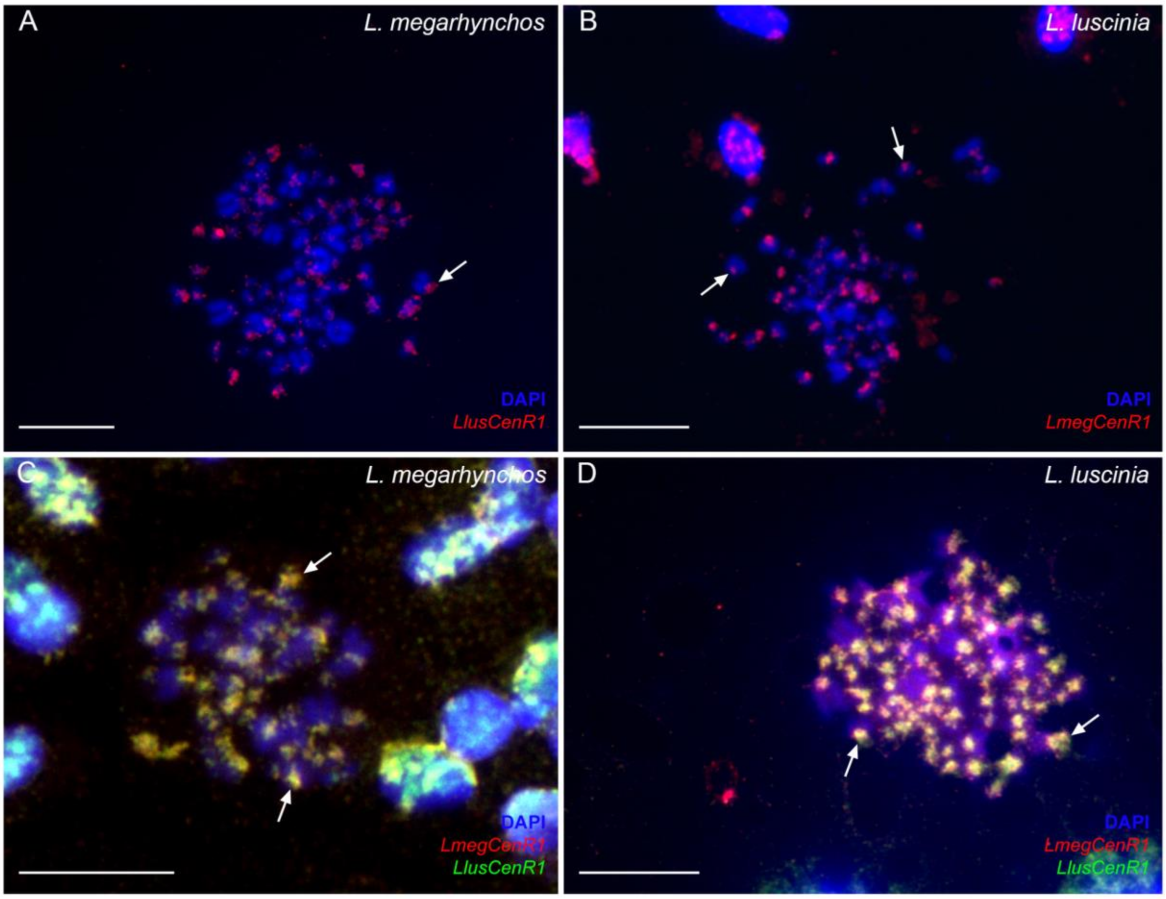


**Supplementary Figure 5.** Cross-species FISH (A, B) and two-coloured FISH (C, D) of CenR1 probes on *L. megarhynchos* (A, C) and *L. luscinia* (B, D) mitotic chromosomes. *LlusCenR1* repeat localizes in centromeric regions of *L. megarhynchos* chromosomes (A) and *LmegCenR1* repeat localizes in centromeric regions of *L. luscinia* chromosomes (B) (indicated by arrows). *LlusCenR1* and *LmegCenR1* repeats colocalize in the centromeric regions of all chromosomes in both species (C, D) (indicated by arrows). Chromatin stained with DAPI. Scale bar = 10 µm.
