## Supplementary tables for "Germline-restricted chromosome of songbirds has different centromere compared to regular chromosomes"

**The document includes:**


Supplementary Table 1
Supplementary Table 2

**Supplementary Table 1**. RepeatExplorer2/TAREAN output for five tandem repeats identified using 200 000 paired-end reads from *Luscinia megarhynchos*

Putative satellites (high confidence)

| Cluster | Proportion[%] | Proportion adjusted[%] | Number of reads | Satellite probability | Consensus length | Consensus | Connected component index C | Pair completeness index P | TAREAN k-mer coverage | \|V\| | \|E\| | PBS score |
|---|---|---|---|---|---|---|---|---|---|---|---|---|
| CL1 | 6.40 | 6.40 | 14415 | 0.986 | 436 | TCTGCACGACGAATAACTATGTTGTAGCAAAGCTGGTTCTGGCATATCCTGCTTCATACCACTAGTCCCTGACAACCGGT<br>ATTCTGTGCAGAGCACCTCAGTGCAGTGTGGTTTCCCTAATCGCGGTTTGTAGCAAGAAGCGATTTCTGTCTAAGGGAGG<br>AGAAGTGCCGTCTACCCCTCTCACAGAACACTCTGCTGGGAAGTTCTGCTACTGCCAAAACACGCCTGGCATGTGCACTT<br>TTCTTCTAAAACCTCTCTACCATAGCAATATTCTCTGCTTCTTGAAAGCATACCATACACTCTTTCCACTATGCATAGCG<br>AAGAGCACTGCAGTGCACCTAAAGCTAAAGCGAAAGTGTGCACAGCACACCACAGATCTCACTTTGCACTACACCTAGAA<br>ATGGCTTCTCTGCATCACTGATATCTAGCACAGAAA | 0.996 | 0.966 | 0.842 | 10123 | 19999088 | 0.000 |
| CL9 | 0.19 | 0.19 | 437 | 0.846 | 495 | ATGTTTTCCCCAATAATGCTTGGAATTTTGACCGGTTTGGGCTGTTTCAGTGTCGATTTTAGGACTATTTCTGGCCATTT<br>TCATATGGATTTTGAGGGGATTTTTTGTTTCTTAGTGCCAATTTGTTAGGCTGATTTTCCCTGGATTTTTCCTGATGCTT<br>TCCCCAGCAGTCCTTGGAGTTTTGGCCGAGTTTACGTGTTCTAGTATGGATTTCAGGAGTCTTTGGGCCATTGTTCGTGT<br>CATTTCCACACTTTTTGGGCCATTTTATGTCTCATTTCCTCTGGATGTTTTCTCCGCTCATTTCCCCAGTTTTTCCTCAG<br>AAATTCCAGGAATCTCACTGGATTTTGGGGCCATGTTTGGCTTTTTAGTGCAGATGTTAGGTGTTTTTTCTGGCCATTAG<br>GGTGCAGTTTCTGGGGGATTTCTGTGTTTCTTAGTGGGAATTTTTAGGCATTTTCCTGGGTTGCTAAGGTTCAATTTATG<br>CTGGATTTTTTCCTG | 0.947 | 0.917 | 0.304 | 437 | 7843 | 0.279 |
| CL12 | 0.12 | 0.12 | 273 | 0.737 | 1309 | ATGAATGTGGGGGCGGGATTTTGTTTCACAGGTTAACGGGCCACTATTTGAGTCCAGTGCCCTGGGCGTGAGAATTAAGA<br>CCCCTGCTAAAATGAGAAACGGATGAATATTTCTCCCAGATGCAAGGTGATATCATTTAGGTCTATGTCGGCCTGCAGAG<br>TGACAAAGGCCTTCTCCAAGCTGGTATCGGGAAACAGGGAGGGTGAGTACTTTTTCAAAGAGAAATAAGGCCATTATTTG<br>TGTCCAGGGCCCTGTGGCTGACACCGGGCTTCTGCATGCTGAAATTCTGAACGGATCAGCATTTCTCCCAGACACAAACT<br>GACACCACTTAGAACCCTGGCTGCTTTTTGAGTGCCAAAAGCCTCCTGTGAATAGACATTACAAAAGTGGTGAAGAGAGC<br>TGTATTTGACAGAGAAATGGGCTCGTTTCTGTCCATTGCCCAGTGAATGAAACGGGGCTTCTGCCTGCTAAAATTCCAAA<br>CAGTTCAGGATGTCTCACACACAATCTGGCAACACTTAGAACCCTGGCTGTTCTTTGAATTCCAAACACCCGCTCTAAAC<br>TGACATTCCAAAAGTGGTGAAGTGAGCTGTATTTCACAGAGAAATAGGCCCCTGTTTGTGTCCATTACCCAATGAATGAA<br>GTGAGGCTTCTGCCTGCTAAAACTCCAAACTGTTTGGAAATCTTGCAGATAGAAACGGGCACCATTAGAACCCTGGCTGT<br>TCTTTGGCTGACATCACCCAGTCTAAACTGACCTTATGAATGTGGGGACGGGGTTTTTGTTTCGCAGGTGACAGGGCCAC<br>TATTCATGTCCAGTGCCCCGGGCCTGAGAATTAAGTCCGCCTGCTAAAATGAGAAACAAATCAATATTTCTCCCAGATGC<br>AAGCTGCCATCACTTACAGCTATGTCTGCCTGCAGAGTGACAAGGGCCTCCTCCAGACTGGTATCGGGAAACTGGGAGGG<br>TGAGTATTTTTTTCAAAGTGAAAAGAGCCAATGTTGTTATCCATGGCCCTGTGCCTGAGACTGGGATTCTACACGGTAAA<br>ATTCTAAACCGAACAGTATTTCTCCCAGACTAAAGCTGCCACCACTTATAGCCCTGGAGGTCTATCCTGTGCCAAAAGCC<br>TCCTCCAAATCGACATTAAAGAAGTGGCGAAGTGAGCTCTATTTCACAGAGAAATAGGTCCCTGTGCCTGCGACTGGGCC<br>CTGTGCCTGCTAAAATTCCAAACTCTTAGGGAAGTCTCGCAGATACAGACTGGCACCATTAAAACCCTGGCTGTTCTTTG<br>AATGACATCAACCAGTCTTAACTGACCTT | 0.941 | 0.923 | 0.838 | 273 | 4564 | 0.478 |

Putative satellites (low confidence)

| Cluster | Proportion[%] | Proportion adjusted[%] | Number of reads | Satellite probability | Consensus length | Consensus | Connected component index C | Pair completeness index P | TAREAN k-mer coverage | \|V\| | \|E\| | PBS score |
|---|---|---|---|---|---|---|---|---|---|---|---|---|
| CL6 | 0.320 | 0.320 | 717 | 0.035 | 2643 | CGGAGCAGTCTCTGGTGGGGTCTCCGCGCCCTTCGTGGCCGGGCAGGCGGCACAGAGCGGATCTCCGCCCGCTCCGTGGG<br>TCTCCTCCCGGTGCGCTGCTCGGGGCTCGTCTCCCAGGGACCCCTCCTCCTCTTTCCGCCGCCGTCTCTGCTGAGGGCCC<br>GAGCCAGCGGGGCTCTTTATATACCCAGACCAAGCCCCGCCCCTTTCAGGCACAGCCAATAGGAGCCCAGGAACAAGCGT<br>GGGGGCGGAGCACCCAGCGTTGCTCATTGTGATGTCACAAAGAGGAGGCGTGGCCACAGCGTGACCGAGCGTTCTGATTG<br>GTCGGAAGGAGAAAGAGCAGAAAAGGGGCGTGGCTCAGGCCCGGGGTGGGAGGGGCAGGTGTGAGGGCGGGGTGGGGGCG<br>GGGCCAGGGTCGCACCGGCATTTTGGGAGCAGGTCCGGGCGCTCCCGGTCCCTCCACAGCTCTCCTGGGCCCCTTCAGGC<br>ACTGCAAGAAGCAGAGGAAGGGGTCTCTGGGCGTCCCAGCCCCATTCCAGGAGTCGACAAGTGTCGACAGCGTGACGAGA<br>AGTTGCAACACTTATGGCTACAAGGACTTTTAAGGATTTATTCACCAATAACACACTGCCAACAAAGAGATTTATGTTTT<br>TAATGCAATACTAAAATTTTGTCTAATTGCAGTGTGAACTTTCTTAGCCAATCCTGTTCTGGCACACAGACCTAGTCTAA<br>GACTTTTTGTCACATCTGCGACTAATTTTATTCTTCTTTTTATCATAAAATCTACAGCTAAACTTTTCTCACCTAAATAC<br>ATCTCTGCTTTAATCTACATTTTTGCTTCTAGCACCTAAGTTGGAAGCATTTCCTGAGGCCTCCGGTCAAATCCTGTGTT<br>CTTTTCTAGACTTTGATTGACAGGCCCAATGTCGAGAGAATTCCTTGCATTTTGAATTCCAACAGAATCTCACAGACCCG<br>AGGCTCTCTTGTGACACAGTAGGGCCTGGTGGAACCAAGGGCAGCGTTGTGACACCCCAGAACCTCACGGAACCGAGTGT<br>CAGTCGAGACACAGCGGGGCCTGGGGAAAGGAAATCTCACTGCCAAAGCATTTCTGGCTGATAAAAGTTTGATTTGGGCC<br>AACACCCAGGGATGCTCGAGGGTCTCCATCCCTCTGGGCCTGGGAAATGCTCGAGGTGGCCGGGGGGTTTCTGTCCCTCT<br>CCCTTGGGGGTGGGGTCTTCATCCCTCTGGAGACACCGCAGGGTCTCTGTCCCTCTCACCCGAGGATGAGCGCCCCGAGC<br>ATCCCCCGGTGTGACACGGCCAGAGACCTCCAGCCCTGCCATTCCCCTCAGGTGAGGCCCGAAACCCCCCAGACCGCCCA<br>GGGAGGAGGGGAACAAAGACATCCCTCGGCCCACCGCGCTCCCTGCCCGGCTGAGGGTCTCTGTCTTCCTCTTCATGGGG<br>CGGGGGGGGCTTTGAGCTTGGTTTGGTTGGCGAGGGACTCGTTGCGAGAGTAAAGACCAGGCACAGCACTTGATGTGATT<br>TTTGCCAATTTTATTTTGGCCTTCGATTTTTTGAAGCGGGACGATGGCCAGACCCCTCCAATTTGGGGGGCCGGCGCGGT<br>CCCTCGCTGCCCCGGTGCTTGTCTGTCCTTAGCTTTGTTGAGCCGGGGTGGTGTCCGACCCCCTCCGATTTTGGGAGGCC<br>GCTGTGCTCTCCATCCACCTTACCTTTGGCAGGCTGTGCTGTCTTTGTCACCTTTTGGCGGGGAGGGGAAAGCGCTGTCG<br>GTGTCCGTCATCCTTGCCCCCGGGGGGAGGCTCACCCCCTGTCCTGGGGTGACGTTGCAGTGCTTTATATCCACAATCGT<br>GTGCTGCTCATGCCTGGAGGGTTTGTTTTGTCTAGCTAGACAGCGCCCCTCCCCATCGGTGAGCTGCTGCCGCTGTGCTG<br>CTTGCAAGCTGCTGCCCCCTCTCCCCCCCTGCGCGGCTCGGCCGCTTTTAGCTGCTTCTGCTTGCTTAGCTGCTGCTTCT<br>GCCCCCCACTTTGCTTAGGCTGCTTCGCTTTATTTTTCCTTTCCTTATTAGCTAGAAATACCACGTGAACCTGATCCTGG<br>ACGACCTGGAACATCAAGAAACGCCGGGCGCCTGCTCGTGAAAAGAAATACGCGTTTCTGCCCTCCCCGGCACGGGAGAT<br>GTCCTGAGGCTGTACATCCACTCGGACGGGTGAAGTTTGCTGCGTCAACCTTGGCTCTTTTTTTCTTGCGTTGGAGAGAG<br>ATTTTTCTGTGTGTTTGCTAAATAAACAAGTGTTTTCGCCTTTTTCTCTGAGGAAAATCTTCCCGAAGCCGGTGGTGGGA<br>GAGGCAAGAGTTTGGAGAGTTTGTTGCTTAAGGGGTTCCTTTTGGAGATTCTTCCCCTAATTTGCCCTAAACCAGGACAC<br>GCCTGTATCTGCCCTCAGCCCAGGGCGATGCTCTGTCCGGCTCCGTCATTGCCCCGGGGAGATGCTCTGTCGGTGTCCGC<br>CCTCCTCGCCCCGAGGGAGGAGCGGCTTGTCTCGCCGTGCCCATCCTTTCTGTGTGGGGATGGTGGGGGTGGTCAGGACA<br>TGTCCGCCTGCTTCCCATTACTGGGAGGGGTCTGGGTGGCCTCCGTGCTTACCTTGGTGTGGGTGGGGGTGGTCTCCATG<br>GAG | 0.752 | 0.862 | 0.443 | 717 | 7393 | 0.387 |
| CL20 | 0.063 | 0.063 | 143 | 0.258 | 1883 | TTTTTTGGTTGCTGCTTTGTGAGGAGGAGAAGCCCTGCTCATTTTCTCTTGATGAAGAAAGCAAAAGCATTTTGCTCAAA<br>ATTTCTTCCCCTTTGAAGAAAAGGATGAGATTCTGTTAGTGTTTTCATTTTCCATGTCTGCAGCAGCAACAGCTGGGCAT<br>TTCCTAATTTTCCATTCTAGTGTTTTAAGGAACTCAAAGGTCAGAATTCAGACACAACCTTGCTTGCCGACAGTATCCAT<br>TCTGGAATATTGAATTCAAAGCCACATGCAGTGCTGTTGTAGCAGCCCATCTTAATTTAGTTGGAGTGACTTTGAATCCC<br>ATTGCTATGACCGTTTATGGTTCTGCCTCAGTAGATCAGGTTGTAGAGCTGAATAGAACTGGGATTCCAAATGATAGATA<br>ATGGCTCATTCCTCATGCTTCTCTCTGTCCACAGCGCCCAGAAGCAGCCCCAGACGATGCAGTGACCCTGTCAGCTGCTG<br>GAGATGAAGCCATAGAGGAGGAAAATGACAACAAGCCTCCAGCTATGGTCCCAGCAGAAGCTGAATATTCGGAGATGGTA<br>AGTGCCACTTCTGGCTGGTGCTGTCTTTATTGCAAGAGAGCGCTGCAACTTTCTGAGCAGCCAGCCCTTGCCGTTGGTGG<br>CTGAGGGTGCTGAGTGCGGGAGTCCTGCCAGGAGCTAGCACTCCCTCTTGGCAGTGACAGGTGGAAAATGGGCTTAAATC<br>TGGCATGATTTGGCCGCAGTCTGCTGGGATTGTGATAAGGGCCAGTCTGAAGCACGAAGGCTCTTTGGCCCTAGAGGAGG<br>GAGTGTGTCAATGACACAGTTGTGATGGACACGTAGCAGGACACTGTGGTCTGGGGAGACGTGTCCCACGGCAGAGTCCT<br>GCCAAGGAGTTACAAGCACAAAAATGGGTGATCTGAGCCCACAGTCACTTTGGGGTGCTTTGTCCTCACCAGATGCTTTT<br>CTCTACTTGCCTGGATTCCATTAAGAGACACATTGTGGAGTTGCAGCTTGAAGTGGCAGATTTGGTGTCTTTATTGCCAG<br>GTGGTTCTTTTGCCTCTTCGGCCAGAGCAGTCAGTAGCAGAGTGCAGTCTGAGTGCAGCTGTGTTCTGATAAGGTGCAAA<br>GAGATGATTGTGGTTTCCCTGAATGCTTGTGGGCCTACTCACATCTTATGTGTCAAAACAGCGAAATTGCATACTTGTGG<br>ACCAATTTCAGAGGCAAAATCCAGATGCTACATGAATCCCGGGATACTTTGTGTTGCCTCATGTGCCTTGGTAACTCTGT<br>TTAAAAATGCCATCGAAGCCTTCAGAGGGGAAAAGCTGCACAAGCTTGACTGCAGTCTTGTGTTGCTTAAACGAAGAAGG<br>TTTTGGAAATGCGGATGCTGTATCCAAAGTCAAATGTGCAAGTTTGCATCTGGGGAGCTGCCTGAGACAAGGGGAACCAG<br>TGTTGCCCAACAAGAGCAAGTGAGCAAGAGGCAGCGCGTGCCCAGGTGGCCAAGAAGGCCAAGGGCATCGTGGCGTGTGT<br>CAGCAGCTGTGGGGCAGCAGGAGCAGGGCAGTGAGCTGTGCCCTGTGCTGGGCACCGCTGCGGCCGCTGCTCGAGTGCTG<br>TGTCCAGTTGTGGGCCCCTGTGGAGCAGAAAGACATTGAGGGGCTGGAGCGTGTCCAGAGAAGGGCAAGGGAGCTGGGGA<br>GGGGTGGGGGAACAAGCTCAGTGAGGAGGCGCAGAGGGATCTTTAGCCTGGGGAGAGGGAGGCTCATGACGGACATTGCA<br>GGCTCAATTCCAGAAATGCCTACTTGACAGTGCTCCCCACAGGGTCTGTTTCTGTGACAAGAATGCCCTCAGTGCATCCA<br>GATGCTTTAAGCATTGGAGTCGGTGACACTTTGATCGTTTGGT | 0.916 | 0.810 | 0.834 | 143 | 729 | 0.739 |

**Supplementary Table 1**. RepeatExplorer2/TAREAN output for eight tandem repeats identified using 200 000 paired-end reads from *Luscinia luscinia*

Putative satellites (high confidence)

| Cluster | Proportion[%] | Proportion adjusted[%] | Number of reads | Satellite probability | Consensus length | Consensus | Connected component index C | Pair completeness index P | TAREAN k-mer coverage | \|V\| | \|E\| | PBS score |
|---|---|---|---|---|---|---|---|---|---|---|---|---|
| CL1 | 4.000 | 4.000 | 14560 | 0.993 | 436 | ACATAGTTATTCGTCGTGCAGATTTCTGTGCTAGATATCAGTGATGCAGAGAAGCCATTTCTAGGTGTAGTGCAAAGTGA<br>GATCTGTGGTGTGCTGTGCACACTTTCGCTTTAGCTTTAGGTGCACTGCAGTGCTCTTCGCTATGCATAGTGGAAAGAGT<br>GTATGGTATGCTTTCAAGAAGCAGAGAATATTGCTATGGTAGAGAGGTTTTAGAAGAAAAGTGCACATGCCAGGCGTGTT<br>TTGGCAGTAGCAGAACTTCCCAGCAGAGTGTTCTGTGAGAGGGGTAGACGGCACTTCACCTCCCTCAGCCTGAAATCGCT<br>TCTTGCTACAAACCGCCATTAAGGAAACCACACTGCACTGAGGTGCTCTGCACAGAATACCGGTTGTCAGGGACTAGTGG<br>TATGAAGCAGGATATGCCAGAACCAGCTTAGCTACA | 0.995 | 0.959 | 0.866 | 10004 | 19979821 | 0.000 |
| CL6 | 0.560 | 0.560 | 2023 | 0.955 | 467 | CAAAATCTCCAGGAATCTGTACAAAAACCATCTACAATCTGCCAAAATACCAAGGATCGATGGGATAGCATGAGGAAAAA<br>TCCAGGGAAAATCAGCCTACCAAATTGGCACTAAGAAACAAAAAATCCCCTCAAAATCCATATGAAAATGGCCAGAAATA<br>GTCCTAAAATCGACACTGAAACAGCCCAAACCGGTCAAAATTCCAAGCATTATTGGGGAAAACATCAGGAAAAAATCCAG<br>CATAAATTGAACCTTAGCAACCCAGGAAATTGCCTAAAAATTCCCACTAAGAAACACAAAAATCCCCCAGAAATTGCACA<br>AAGAATGGCCAGAAAAACTCCTAATATCTGGCCCAAAACCCAGGGAGTGACCTGGACTTTCTAAGGGAAAAAAGTGGAAA<br>AATAACTGGGCAAACTTCCAAAGGAAATGAGACTTTAAATGGCCCAAAATCCCCCCAAAAAATGGCC | 0.977 | 0.936 | 0.383 | 2023 | 233374 | 0.000 |
| CL17 | 0.140 | 0.140 | 520 | 0.956 | 1994 | TGGGTGGTGGCCTGGCGCGAGGGTCTCGATGTGGGGCTGGTGGGTGGTGGTTCTTGTTGGGAGCGGGAGCATTATCAGGC<br>CTGAGTGCGTCGTCATCCGTTGGCAGAAGCCGAGGGACGACGGGGCAAGAGCAAGCTGGATGAGTGAGCTTCGAGGTCAA<br>GAGTGGAACGGCGAGGCAAGAGCGAGCTTGGAGCCGAAGAAGGGAGCACAAGAGGCCCTCTGGGCAAAGGGCCTTAATGG<br>CACGGCCAGGGCACCTTTAAGGTGCCGTGGCTGACTTTGCTTGTGCTGGGCGACTTCCCTGCGTGCAGGGTTGCGTCTTT<br>GTTGTGCTTGGCATCTTGCCATTGTGCTCCTTTGGCTGCATGGAGAGAGGGGCAACCCTGTAACGAGGGTGAGCGCTCCT<br>CTCTCCACACCTGAGGGGGGAAACGGAGCTCCCCTAGCCCCCAGCCCGGCCGCCCTGTAAGCAGGGCAGGGCTTTTCCCC<br>AGCCGTGGCAGCTTTCTGCCTGTTGCTGGCCCCTGGCCAGCGTGACCCCCTGTACTCGGGGCGTGTCACCGCCTTCGTCC<br>CCGTAGCAGCCCTGAAGCGCCTCAGCGCGTCGACGCCCCTGTGCCCTCCCCACGGTGCAGTGCGGTGCGGGTCTGATGCT<br>TTTGTCTCCCACGAAGCAGTAGGGATGGCTCCCTTGAGCCCTTTGGCTGCACGGTACAGGCTCCCTCTTCTGGGACTGTC<br>TTCCCCTCCCTCCCTCCCTCCCTTGTCTTTGCATTAGGCGGTGTGAATGAGCCGAGTGCGTTTGTGCTTTAGCCCCCCTT<br>TAGAAATAGAGCGGTGTAGCTTAGGCTTCCCAGGAGAGGAAGAGGAGCTCCGTGCAGCCACAAGCCAGTCCCCAGAATGC<br>TTGCGTGGACAAAGTGTCTGGCCTCGTCCCCTGCTTCCTGAGGACGGTGCGAAGCCGCTTAACGGTAGCGTGGGCAACGT<br>GACCCGCCTGTAGCCGGGGCCTGGGTCTGGGCCTATGGTGCCCTTTGTGCTGGTGAACTGGCAGCGTGCGGGCCTCTTGC<br>TCTGGGCACGTGTCAGGTACTTGCTTGTCCCCTTGTCCCCAGGAGCAGCCCCAGGTCCCTGTAAGCAGGGCTGGGGAGGC<br>TGCTTTAGCCGGGTAACAGGAACCGCTGCGCCCGGGCCTTTGCCAGGGTGCAGGTTGAAGAAGCGCTGGTGCCACCTCGG<br>CGGCAGCCTCGTTGCGGAGGCTGGAGGGGCTTCTGCAGACGGCCATGTTGCCGTGCTGCGCGGCGGCCGGGCTGCGGCCG<br>GCCCCTAATTGGGCGCGGGCCGGAGCCCGTGCTCCCTGGCGCTTGTTGGGGGGTGCCGGGCTGCGGGGGGCGGCCCCGTA<br>AGCGGGGTTGTGTGCGCCCCCCGGTCTCGGCGGGCGGTTGGAGGGACCACCCCCTGTAATCAGGCGGGTGTCCCTGGCCC<br>TGGCCTTTCAGAGGCGTGGGGCCAGGAGTGGTGGGGCCGGGCGTGTGCCCTCCCTGTCGCCAGGGCTGGGCTCCCGGCCC<br>CTGGGGCCCTTGGGCTTTAGGCCGTGAGTTGGGCTGCTCGACTGTGGAGTCTGGTTCCTGTTGCCGTGGGGTGAAGGCCG<br>AGCTCCGTGGCTCCCTGGGTGTGGACCGGCCTGTAAGCAGACTGTGGGTCTGGGCGCCTTTGTGCTCCCTTCAGCTGGCA<br>GCCTGCGTTGAGCGCCCTCCCCGTTAGTGGGGCGGGGGCCCGAATTGGAGAGAAGGAGCCTCCATAGGCCTTGGGACCAG<br>GGCCGCCCTGTAAGTCGGGGCCGGTGCTCTGGAGTCCCGGGCACTTTGCAGTGCAGGCAGCGGCTTGAGCCCCCTGTAAG<br>CAGGGCTGCAGCCTCTTGTGCGCTGCTTTGCTTTGTGCCCAGCTTCTTTTGCCCAGAGGGCCTGGCTCCCAGCACGGCGT<br>GGGACGTTGGCGGACGTCGAGGTGGTGAGTCAAGCTGGAGCTGGAGGCTTTTGGGCGGCTGCGGGCGGGTCCTG | 0.979 | 0.912 | 0.849 | 520 | 11363 | 0.066 |
| CL49 | 0.023 | 0.023 | 85 | 0.741 | 1181 | CTCCAAGAGCCTTCAGCCCTGCCAGAGCCTTGGTGAGAGCTGCCAGAACGCAGTGGGAGGCAGCCCTTGGGACAGGCATC<br>CCTGCTCCCTCAGAGTCACGGCTATGGGACAAACACCCTCATCCCTGAGGCTGCAGATGGGCAGGAAAAGCCCAAAAGCA<br>AAGCTGGGAGTGCAGCCCACACGAGTAAGGCAATGCCAGAAGAATGTGTCCCAGGAGCAAATATGCTGGGGCTGCCAATT<br>GCCAGTGCATGGCTCTGAGGAAAATGCAGTTGTGGCATGGGAGAAGGGAGAGCCTTGGTGTGGTGCCGGACAACGATCCT<br>CCAGCAGCAAAGGCAGCAGCGTTCCCATGTGCAGCTGAGTCTCTGGAGCAGGCAGTGCAGGAGCAGGAAGAGCAAGATCT<br>CCACAACTGGATGACTTCTGGCAAGAGTCTCCATAGCTCCCGGATCCTTTGTGATAGCTGCCGGACTGGAGAGAGAGAGG<br>AAGCCCTTTGGACAGGACTGCCTGCACCCTCAGGTCCGTGGGCATCTGTAAAGTAGGAATGATTCACATACTCAGTCTTC<br>ATACATGCAGGAGTCCTTCTTACAAATGTACATAGATTTAATAACTTGCTGGCTATCAGGTTAGAATAGCTTGAGCCTGG<br>ATAAGCAATAGTATAAATGCTCAGCTGTCAGCTAAGCAATAACATACATGGTCAATCCCCTAGTAGCTACATCAGATTTT<br>TTTTCTTAAGACAAGTGCTTGTAGAACAATTCCATCTTCAAACGACCTTACTTCTGCTACTTAGTCCATTTTAGAACCCT<br>TTTTAGAAATCACATGCCTTAACTGAATAGCGTTCCTTGTTCTCTTATTCCAAATTGTCTTTGTCTTTTTTCCCCACAGG<br>TATCGGCAGAACCTGCTCATCCCTGAGCATGCACAATTGCCAGGGAAATCCAGAAAGCAAACCTGGAGGCCAAGGCCGTT<br>CCAGAAGGTGTTTCCTGAAGCCAGACACACGGCAGCAATTTCCAGAGCTGTCACTTCCTCGGCCACAGCGTTACGCATGC<br>TCTGGCTGTGCCGCAGGACAAGAGCCTTGGTGTTGTCCTGGAACAGGTTCCTGCAGAAGAAAAGGTGGTGTTCCTGTGTG<br>CAGCGAACCCTTTGGCATCAGCAGTAGAGGAGCACAATGCTGAAGATCTCCAGCCTTTTTT | 0.953 | 0.889 | 0.880 | 85 | 505 | 0.442 |

Putative satellites (low confidence)

| Cluster | Proportion[%] | Proportion adjusted[%] | Number of reads | Satellite probability | Consensus length | Consensus | Connected component index C | Pair completeness index P | TAREAN k-mer coverage | \|V\| | \|E\| | PBS score |
|---|---|---|---|---|---|---|---|---|---|---|---|---|
| CL23 | 0.067 | 0.067 | 245 | 0.3930 | 1881 | ACCAAACCATCAAAGTGTCACCGACTCCAATGCTTAAAGCATCTGGATGCACTGAGGGCATTCTTGTCACAGAAACAGAC<br>CCTCTGGGGAGCACGGTCAAGTAGGTATTTCTGGAATTGTGCCTGCAATGTCCATCCTGCGTCTCCCTCTCCCAAGGCTA<br>AAGATCCCTCTGCGCCTCCTCACTGAGCTTGCTCCCCACCCCTCCCCAGCTCCCTTGCCCTTCTCCAGACACCATCAACC<br>CCTCAATGGCTTTCTGCTCCACAGGGGCCCACAACTGGACACAGCACTCGAGCAGCGGCCGCAGCGGTGCCCAGCACAGG<br>GCACAGCTCACTGCCCTGCTCCTGCTGCCCCACTGCTGCTGACACACGCCACGATGCCCTTGGCCTTCTTGGCCACCTGG<br>GCACGCGCTGCCTCTTGCTCACTTGCTCTTGTTGGGCACCACTGGTTCCCCTTGTCTCAGGCAGCTTCCCAGACGCAAAC<br>TTGCACATTTGACTTTGGATACAGCATCCGCATTTCCAAAACCTTCTTGGTTTTAGCAACACAAGACTGCAATCAAGCTT<br>GTGCTGCTTTTCCCCCTCTGCAGGCTTCCACGGCATTTTTAAACAGTTACCAAGGCACATGAGGCAACACAAAGTATCTC<br>GGGATTCATGTAGCATCTGGATTTTCTCTCTGAAATTGGTTTACAGGTATGCAGTTTGGCTGTTTTAACACATGAGATGT<br>GAGTAGGCCCACAAGGATTCATGGACACCACAATCATCTCTTTGCACCTTATCAGAACACAGCTGCACTCACACTGCACT<br>CTGCTACTGACCGCTCTGCCCGAAAAGGCAAAAGAACCACCTGGCAATAAAGACACCAAATCTGCCACTTCAAGTTGCAA<br>CTCCACAATGTGTCTCTTAAAGGAATCCAGGCAAGTATAGAAAAGCATCTGGTAAGGATAAAGCACCCCAAAGTGACTGT<br>GGGCTCAGATCACCCATTTTTGTGCTTGTACCTCCTTGGCAGGACTCTGCAGTGGGACACGTCTCCCCAGACCACAGTGT<br>CCTGCTACGTGTCCATCACAACTGTGTCATTGACACACTCCCTCCTCTAGGGCCAGGGAGCCTTCGTGCTTCAGACTGGC<br>CCTTATCACAATCCCAGCAGACTGCGGCCAAATCATGCCAGATTTAAGCCCGTTTTCCACCTGTCACTGCCAAGAGGGAG<br>TGCTAGCTCCTGGCAGGACTCCTGCACTCAGCACCCTCAGCCACCAACGGCAAGGGCTGGCTGCTCAGAAAGTTGCAGCG<br>CTCTCTTGCAAGAAAGACAGCACCAGCCAGAAGTGGCACTTACCGTCCCCGAATATTCAGCTTCTCCTGGGACCATAGCT<br>GGAGGCTTGTTGTCATTTTCCTCCTCTATGGCTTCATCTCCAGCAGCTGACAGGGTCACTGCATCGTCTGGGGCTGCTTC<br>TGGGCACTGTGGACAGAGAGAAGCATGAGGGATGAGCCATTATCTATCATTTGGGATCCCAGTTCTATTCAGCTCTACAA<br>CCTGATCTACTGAGGCAGAACCATAAACGGTCAGAGCAATGGGATTCAAAGTCACGCCAACGAAATTAAGATGGACTGCT<br>ACAACAGCACTGCATGTGGCTTTGAATTCAATATTCCAGAATGGATACTGTCAGCAAGCAAGGTTCTGCCTGAATACTGA<br>CCTTTGAGTTCCTTAAAGCTCTAAAATGGAAAATTAGGAAATGCCCAGCTGTTGCTGCTGCAGACATGGAAAATGAAAAC<br>ACTAACAGAATCTCATCCTTTTCTTCAAAAGGGAAGAAATTTTGAGCAAAATGCTTTTGCTTTCTTCATCAAGAGAAAAT<br>GAGCAAGGCTTCTCCTTCTCACAATGAAGCCACGGAAAAAA | 0.922 | 0.899 | 0.848 | 245 | 2277 | 0.686 |
| CL28 | 0.054 | 0.054 | 195 | 0.2420 | 63 | CGCCACGGCCGGCCCACCCGTGGGTGGGCGTGCAGAACTGTTTGCGTGTGCCACTGGGTGTGC | 0.954 | 0.789 | 0.630 | 195 | 11447 | 0.000 |
| CL32 | 0.039 | 0.039 | 140 | 0.0147 | 756 | TTTGAATGACATCACCCAGTCTTAACTGACCTTATGAATGTGGGGACGGGATTTTGTTTCACAGGTTAAAGGGCCACTAT<br>TCATGTCCAGTGCCCCGGGCCTGAGAATTAAGTCCGCCTGCTAAAATGAGAAACAAATGAATATTTCTCCCAGATGCAAG<br>GTGCCATCACTTACATCTATGTCTGCCTGCAGAGTGACAAAGGCCTTCTCCAAACTGGTATCGGGAAACTGGGAGGGTGA<br>GTATTTTTTCAAAGAGAAATAAGGCCATTATTTGTGTCCAGGGCCCTGTGGCTGAGACCGGGCTTCTGCATGCTGAAATT<br>CTGAACGGATCAGCATTTCTCCCAGACACAAACTGACACCACTTAGAACCCTGGCTGCTTTTTGAGTGCCAAAAGCCTCC<br>TGTGAATAGACATTACAAAAGTGGTGAAGAGAGCTGTATTTGACAGAGAAATGGGCTCGTTTCTGTCCATTGCCCAGTGA<br>ATGAAACGGGGCTTCTGCCTGCTAAAATTCCAAACAGTTCAGGATGTCTCACACACAATCTGGCAACACTTAGAACCCTG<br>GCTGTTCTTTGAATTCCAAACACCCGCTCTAAACTGACATTCCAAAAGTGGTGAAGTGAGCTGTATTTCACAGAGAAATA<br>GGCCCCTGTTTGTGTCCATTACCCAATGAATGAAGTGAGGCTTCTGCCTGCTAAAACTCCAAACTGTTTGGAAATCTTGC<br>AGATAGAAACGGGCACCATTAGAACCCTGGCTGTTC | 0.714 | 0.818 | 0.466 | 140 | 741 | 1.170 |
| CL59 | 0.018 | 0.018 | 65 | 0.4210 | 80 | CAAAAAATCCCTTCCTAATCTCTGGATTCATGGGGGAGCATTGAAAAAAAGCTGGGATTAAAGGAGCAACTGAGCCAACT | 0.969 | 0.806 | 0.528 | 65 | 603 | 0.000 |
